# Glycosylation-on-a-chip: a flow-based microfluidic system for cell-free glycoprotein biosynthesis

**DOI:** 10.1101/2021.09.24.461663

**Authors:** Alicia K. Aquino, Zachary A. Manzer, Susan Daniel, Matthew P. DeLisa

## Abstract

In recent years, cell-free synthetic glycobiology technologies have emerged that enable production and remodeling of glycoproteins outside the confines of the cell. However, many of these systems combine multiple synthesis steps into one pot where there can be competing reactions and side products that ultimately lead to low yield of the desired product. In this work, we describe a microfluidic platform that integrates cell-free protein synthesis, glycosylation, and purification of a model glycoprotein in separate compartments where each step can be individually optimized. Microfluidics offer advantages such as reaction compartmentalization, tunable residence time, the ability to tether enzymes for reuse, and the potential for continuous manufacturing. Moreover, it affords an opportunity for spatiotemporal control of glycosylation reactions that is difficult to achieve with existing cell-based and cell-free glycosylation systems. In this work, we demonstrate a flow-based glycoprotein synthesis system that promotes enhanced cell-free protein synthesis, efficient protein glycosylation with an immobilized oligosaccharyltransferase, and enrichment of the protein product from cell-free lysate. Overall, this work represents a first-in-kind glycosylation-on-a-chip prototype that could find use as a laboratory tool for mechanistic dissection of the protein glycosylation process as well as a biomanufacturing platform for small batch, decentralized glycoprotein production.

## 1 Introduction

Protein glycosylation is a major posttranslational modification where complex carbohydrates known as glycans are enzymatically added to amino acid sidechains of a protein at specific, regioselective positions. The potential information content encoded in these glycans greatly exceeds that of other biomacromolecules, with distinct glycan structures often playing critical roles in health and disease^1,2^. The attachment of glycans to asparagine residues, known as *N*-linked glycosylation, is the most abundant type of glycosylation and occurs in all domains of life^3^. This mode of glycosylation gives rise to diverse chemical structures that are well known to affect the biological and biophysical properties of a protein^4–7^. Because of these pronounced effects, there is a strong incentive to study glycosylation and leverage the resulting knowledge for the development of glycoengineered proteins with advantageous properties^8–11^.

In eukaryotic *N*-glycosylation, glycans are first assembled by glycosyltransferases (GTs) in the cytosol and endoplasmic reticulum (ER), then transferred *en bloc* to the acceptor protein by an oligosaccharyltransferase (OST) in the endoplasmic reticulum, and finally elaborated to final structures as the protein is trafficked through the secretory pathway^12,13^. Thus, unlike the template-driven biosynthesis of DNA, RNA and proteins, glycan biosynthesis is controlled by the availability, abundance, and specificities of GTs and other enzymes involved in glycan synthesis and catabolism^14^. Because of the complexity of this multi-compartment, enzymatic process, products of natural protein glycosylation pathways are typically heterogeneous mixtures of glycoforms that can be difficult to isolate from the array of intermediate glycoforms and side products. As a step towards producing more homogeneous glycoprotein products, efforts have been made to better understand, control, and expand glycan synthesis in eukaryotic cell-based systems^15–18^. However, an inherent challenge of engineering existing glycosylation networks in eukaryotic cells is that *N-*linked glycosylation is an essential function, so modifications to these networks for the purpose of altering the target glycoprotein product can have adverse effects on the cell. Thus, even with the availability of powerful genome editing tools such as CRISPR-Cas for glycosylation engineering^19^ there are strict limits on the extent of top-down engineering that one can achieve in eukaryotic host cells. As such, there remains a need for alternative methods to produce structurally uniform glycans in sufficient quantities for mechanistic studies and other downstream applications.

To this end, the emerging field of cell-free synthetic glycobiology has helped to expand the glycoengineering toolbox with new methods for synthesizing glycomolecules outside the confines of living cells^20,21^. In these approaches, glycosylation enzymes and substrates are synthesized and assembled *in vitro* to form multistep glycosylation pathways, with the simplest forms involving purified components such that the reaction composition is well-controlled^22^. Alternatively, glycosylation enzymes and substrates can be prepared by cell-free protein synthesis (CFPS) individually^23^ or in a single-pot reaction^24^ to circumvent labor- and time-intensive protein purification steps. The advantages of these and other “open” formats for synthesis of glycans and glycoconjugates include enhanced control over reaction conditions, decoupling of glycosylation and protein synthesis from cell viability, and the ability to use enzymes from any and/or multiple host cells in the same system. Moreover, cell-free biomanufacturing is amenable to real time monitoring, automation, and continuous manufacturing systems.

In the context of CFPS, microfluidics offers a unique opportunity to build scaled-down models of integrated protein production systems in a format that enables precise and tunable spatiotemporal control, usage of small volumes that minimize waste, experimentation on length and time scales similar to those in cells, and in-line process monitoring through real-time, high resolution imaging^25,26^. Indeed, microfluidic systems have been shown to improve CFPS in many ways^27^. In particular, protein yields from microfluidic CFPS systems were measurably increased compared to those of traditional one-pot CFPS reactions as a result of greater heat and mass transfer^28^ and the exchange of reactants and waste products through dialysis membranes^29^ or engineered nanopores^30^. Furthermore, CFPS has been combined with affinity purification in integrated microfluidic systems, enabling efficient protein synthesis and capture^31,32^. With respect to cell-free synthetic glycobiology, there has only been one report describing the use of a microfluidic system in combination with a glycoenzyme^33^. In this seminal work, a digital microfluidics chip was used to merge a droplet containing the soluble GT enzyme D-glucosaminyl 3-*O*-sulfotransferase isoform-1 (3-OST-1) and its adenosine 3′-phosphate 5′-phosphosulfate (PAPS) cofactor with a second droplet containing heparin sulfate (HS) glycans immobilized on magnetic nanoparticles. Following merging of the droplets on-chip, the HS-nanoparticles became enzymatically sulfated as determined by off-chip analysis of the immobilized HS glycans. To our knowledge, however, there have been no reports of microfluidics-based cell-free protein glycosylation.

Here, we developed a first-in-kind microfluidic device for achieving controllable biosynthesis of glycoproteins, which involved reconfiguring a one-pot method for cell-free glycoprotein synthesis (CFGpS)^24^ into a microfluidic architecture. Our prototype involved spatiotemporally separating protein synthesis and protein glycosylation, akin to the subcellular compartmentalization that underlies the biosynthesis of glycoproteins in eukaryotic cells. Specifically, we modeled the cytosol and ER with a modular device that is capable of continuously synthesizing (module 1) and glycosylating (module 2) proteins, after which the post-translationally modified protein products were enriched from the reaction mixture by affinity capture (module 3) (**Fig. 1**). Our results demonstrate that the resulting device was capable of site-specifically glycosylating a model protein, namely superfolder green fluorescent protein (sfGFP), with a bacterial heptasaccharide glycan at a defined C-terminal acceptor site. Importantly, this work represents the first enzymatic glycosylation of a protein substrate in a microfluidic device and a critical first step on the path to building more complex reaction networks for *N*-linked protein glycosylation that more closely mimic the highly coordinated and compartmentalized process in eukaryotic cells.

**Figure 1.**
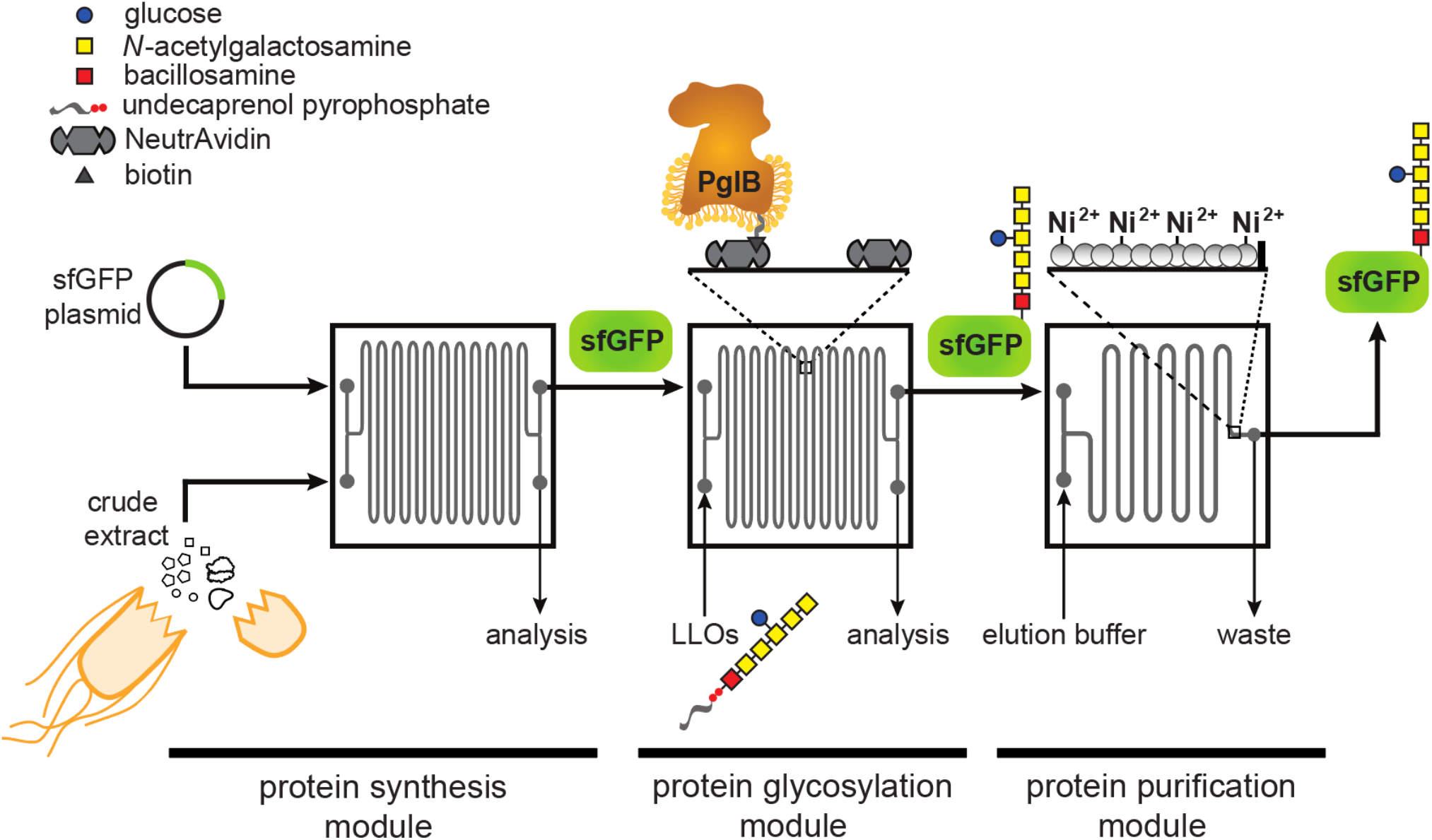
Schematic of glycosylation-on-a-chip system. The microfluidic platform integrates cell-free protein synthesis, glycosylation, and purification. In the first module of the device, one stream containing *E. coli* cell-free extract and a second stream containing plasmid DNA encoding the acceptor protein are combined at the inlet and mixed by diffusion as they travel through the channels. The product of the first chip is then delivered to a second module where it is subjected to an environment enriched with glycosylation machinery. In this case, glycosylation machinery is derived from *C. jejuni* and includes: (i) the OST enzyme, *Cj*PglB, that is tethered to the surface of the device and serves as the conjugating enzyme; and (ii) *Cj*LLOs comprised of undecaprenol-pyrophosphate-linked heptasaccharide from *C. jejuni* as the glycan donor. In the third module, protein product is isolated using immobilized metal affinity capture (IMAC). Depicted is the *C. jejuni* GalNAc_5_(Glc)Bac heptasaccharide with reducing end bacillosamine (Bac; red square) followed by five *N-*acetylgalactosamine residues (GalNAc; yellow squares) and a branching glucose (Glc; blue circle). Structure drawn according to symbol nomenclature for glycans (SNFG; https://www.ncbi.nlm.nih.gov/glycans/snfg.html).

## 2 Materials and Methods

### 2.1 Bacterial strains and plasmids

*E. coli* strain DH5α (lab stock) was used for all molecular biology. *E. coli* strain BL21 Star™ (DE3) (Novagen) was used for expression and purification of sfGFP containing a C-terminal glycosylation tag^34^ and polyhistidine tag (sfGFP^DQNAT-6xHis^), which was used for *in vitro* glycosylation reactions. *E. coli* strain BL21 Star™ (DE3) was also used for expression of the enzyme BirA, which was used for biotinylation of the *Campylobacter jejuni* OST enzyme PglB (*Cj*PglB), and for preparing crude S30 extract. *E. coli* strain CLM24^35^ was used for expression and purification of *Cj*PglB while *E. coli* strain SCM6^36^ was used for preparation of lipid-linked oligosaccharides bearing *C. jejuni* heptasaccharide glycans (*Cj*LLOs).

For cell-free expression of sfGFP^DQNAT-6xHis^, the pJL1-sfGFP^DQNAT-6xHis^ plasmid^24^ was used. Plasmid pTrc99a-BirA (lab stock) was used for expression of the BirA enzyme. Plasmid pSPI01A-*Cj*PglB encoding *Cj*PglB with a C-terminal AviTag was constructed as follows. First, the *Cj*PglB^10xHis^ gene was PCR amplified from plasmid pSN18^37^ and the resulting PCR product was then ligated between the NdeI and EcoRI restriction sites in plasmid pSPI01A^38^, a vector containing the AviTag after the EcoRI cut site. All plasmids were confirmed by DNA sequencing at the Biotechnology Resource Center of the Cornell Institute of Biotechnology.

### 2.2 Protein expression, biotinylation, and purification

Preparation of lysates containing *Cj*PglB with a C-terminal AviTag was performed according to previously published methods^18,24^. Briefly, a colony of *E. coli* CLM24 carrying plasmid pSPI01A-*Cj*PglB was grown overnight in 5 mL of Luria-Bertani (LB) media supplemented with chloramphenicol. Overnight cultures were then subcultured into 1 L of terrific broth (TB; 24 g/L yeast extract, 12 g/L tryptone, 8 mL glycerol, 10% (v/v) 0.72 M K_2_HPO_4_/0.17 M KH_2_PO_4_ buffer) supplemented with chloramphenicol. Cells were grown at 37°C until an optical density at 600 nm (OD_600_) of ∼0.6 and then induced with 100 µM isopropyl β-D-1-thiogalactopyranoisde (IPTG) for 18 h at 16°C. Cells were harvested by centrifugation, after which the pellet was resuspended in Buffer 1 (25 mM TrisHCl, 250mM NaCl, pH 8.5) and lysed using a C5 Emulsiflex homogenizer (Avestin). The lysate was centrifuged to remove cellular debris and the supernatant was ultracentrifuged at 120,000 × g for 1 h at 4°C. The resulting pellet was manually resuspended using a Potter-Elvehjem tissue homogenizer into buffer 2 (25 mM TrisHCl, pH 8.5, 250 mM NaCl, 1% (w/v) n-dodecyl-β-D-maltoside (DDM), and 10% (v/v) glycerol). Once fully resuspended, the solution was rotated at room temperature to facilitate solubilization of the protein and then ultracentrifuged again at 120,000 × g for 1 h at 4°C.

To prepare BirA-containing lysate, BL21(DE3) cells carrying pTrc99a-BirA were grown overnight and then subcultured into 250 mL of LB media supplemented with kanamycin. Upon reaching an OD_600_ of ∼0.6, cells were induced with 100 µM IPTG for 18 h at 30°C. Cells were harvested, resuspended in Buffer 1, lysed by homogenization, and centrifuged to remove cellular debris. To prepare biotinylated *Cj*PglB (*Cj*PglB-biotin), *Cj*PglB-containing lysate was mixed with BirA-containing lysate and 5 mM biotin, 10 mM MgCl_2_, 10 mM ATP, and 1 EDTA-free protease inhibitor cocktail tablet (Thermo Scientific). The mixture was rotated overnight at 4°C to allow time for biotinylation. *Cj*PglB-biotin was then enriched using HisPur Ni-NTA resin (Thermo Scientific) according to manufacturer’s recommendations and the elution fraction was desalted with buffer containing 50 mM HEPES, pH 7.5, 100 mM NaCl, 5% glycerol (v/v), and 0.05% DDM.

To prepare sfGFP^DQNAT-6xHis^, BL21(DE3) cells carrying plasmid pJL1-sfGFP^DQNAT-6xHis^ were grown overnight and subcultured in LB media supplemented with kanamycin. Upon reaching an OD_600_ of ∼0.6, cells were induced with 100 µM IPTG for 18 h at 30°C. Cells were collected, resuspended in buffer containing 50 mM NaH_2_PO_4_, pH 8, 300 mM NaCl and lysed as above. The sfGFP^DQNAT-6xHis^ was purified using HisPur Ni-NTA resin as above. The final protein was desalted using buffer containing 20 mM HEPES, pH 7.5, 500 mM NaCl, 1 mM EDTA.

### 2.3 Solvent extraction of *Cj*LLOs

*Cj*LLOs were prepared by organic solvent-based extraction according to a protocol that was adapted from previous methods^24,39^. Briefly, SCM6 cells carrying plasmid pMW07-pglΔB^40^ were grown overnight in LB media supplemented with chloramphenicol. Cells were then subcultured into 1 L of TB media, grown at 37°C until reaching an OD_600_ of ∼0.7, then induced with a final concentration of 0.2% (w/v) *L*-arabinose for 16 h at 30°C. After induction, cells were harvested by centrifugation, the pellet re-suspended in methanol, and the cells dried for two days at room temperature. After drying, the cells were collected and subsequently suspended in 12 mL 3:2 (v/v) chloroform:methanol, 20 mL water, and 18 mL 10:10:3 (v/v/v) chloroform:methanol:water. After each step, sonication was used to facilitate extraction of LLOs. After the first two sonication steps, centrifugation was used to separate shorter sugars and water-soluble compounds in the supernatant from the pellet. After the final step, centrifugation was used to pellet the cellular debris and the supernatant was collected and dried at room temperature. After drying, the LLOs were resuspended in buffer containing 10 mM HEPES, pH 7.5, and 0.01% DDM and stored at -20°C.

### 2.4 Fabrication of microfluidic devices

Microfluidic masters were made on silicon wafers according to standard photolithography protocols at the Cornell NanoScale Science and Technology Facility. Briefly, SPR220-7.0 photoresist was spun onto silicon wafers and exposed using an ABM Contact Aligner. Wafers were developed using a Microposit MIF 300. Coated wafers were etched to the desired depth using a Unaxis 770 Deep Silicon Etcher, which was confirmed by using a Tencor P10 profilometer. Remaining photoresist was removed via plasma cleaning, and a coating of (1H,1H,2H,2H-perfluorooctyl) trichlorosilane (FOTS) was applied using a MVD-100 to allow for easy removal of polydimethylsiloxane (PDMS). Microfluidic devices were made by pouring degassed PDMS (mixed 1:10 with crosslinker) and curing for 5 h at 60°C. PDMS molds were cleaned with ethanol and MilliQ water, before being dried with nitrogen gas. Final devices were assembled after oxygen plasma cleaning at 700 μm for 25 sec and sealed with a Piranha washed (70/30 (v/v) H_2_SO_4_/ H_2_O_2_ for 10 min) glass coverslip. Devices were placed in a 70°C oven for 10 min to promote bonding of the PDMS to the glass.

### 2.5 Cell-free protein synthesis

S30 crude extracts for CFPS reactions were prepared using a simple sonication-based method^41^. Briefly, BL21(DE3) cells were grown in 1 L of 2xYTPG media (16 g/L tryptone, 10 g/L yeast extract, 5 g/L NaCl, 7 g/L K_2_HPO_4_, 3 g/L KH_2_PO_4_, 20 g/L glucose) and harvested upon reaching an OD_600_ of ∼3.0. Cell mass was washed three times in Buffer A (10 mM Tris-acetate, pH 8.2, 14 mM magnesium acetate, 60 mM potassium glutamate and 2 mM dithiothreitol) then resuspended in a ratio of 1 mL of Buffer A to 1 g wet cell mass. The resuspended cells were sonicated with an optimal energy input (reported based on the volume obtained after resuspending cells) and centrifuged at 30,000 × g to obtain S30 extract, and the supernatant stored at -80°C. No run-off reaction was needed for the BL21(DE3) extract.

CFPS reactions consisted of a mixture of components at a final concentration of 13 ng/μL plasmid DNA, 40% (v/v) S30 crude extract, 1.2 mM adenosine triphosphate (ATP), 0.85 mM guanosine triphosphate (GTP), 0.85 mM uridine triphosphate (UTP), 0.85 mM cytidine triphosphate (CTP), 34 μg/mL L-5-formyl-5, 6, 7, 8-tetrahydrofolic acid (folinic acid); 170 μg/mL of *E. coli* tRNA mixture, 130 mM potassium glutamate, 10 mM ammonium glutamate, 12 mM magnesium glutamate, 2 mM each of 20 amino acids, 0.33 mM nicotinamide adenine dinucleotide (NAD), 0.27 mM coenzyme-A (CoA), 1.5 mM spermidine, 1 mM putrescine, 4 mM sodium oxalate, 33 mM phosphoenolpyruvate (PEP), 100 μg/mL T7 RNA polymerase.

For CFPS in a microcentrifuge tube, 15-μL reactions were conducted in 1.5-mL microtubes in a 30°C incubator. For CFPS on-chip batch reactions, CFPS reaction mixtures were manually inserted into the microfluidic device using a syringe and incubated at 30°C in a moist environment to prevent evaporation. For CFPS on-chip reactions with continuous flow, two mixtures were prepared— one containing S30 crude extract and T7 RNA polymerase and the other containing the rest of the CFPS components— that when combined contained all components diluted to the final concentrations of a standard CFPS reaction. These mixtures were flown into the microfluidic device using a syringe pump where the reactants had a total residence time in each chip of 30 min.

### 2.6 Preparation of functionalized surfaces

Silane-PEG5000-biotin (Nanocs, Inc.) was dissolved in 95% (w/w) ethanol in water according to the manufacturer’s recommendations. The solution was manually pushed into the microfluidic devices and left to react for 2 h at room temperature. Devices were flushed with 100 μL of MilliQ water and then PBS at a flowrate of 10 μL/min. A solution of 100 µg/mL NeutrAvidin (Thermo Scientific) was then introduced and left to bind to the surface for 1 h. For loading of purified *Cj*PglB-biotin, the devices were rinsed with PBS and then buffer containing 50 mM HEPES, 100 mM NaCl, 5% (v/v) glycerol, and 0.01% (w/v) DDM at pH 7.5. The purified *Cj*PglB-biotin was then introduced into the device and allowed to bind overnight at 4°C; unbound enzyme was rinsed away before use.

### 2.7 On- and off-chip glycosylation

For off-chip *in vitro* glycosylation (IVG) reactions, mixtures consisted of components at a final concentration of 17 μg/mL of purified sfGFP^DQNAT-6xHis^, 170 μg/mL solvent-extracted *Cj*LLOs, 10 mM MnCl_2_, and 0.1% (w/v%) DDM. For microcentrifuge tube reactions, IVG reaction mixtures were supplemented with 170 μg/mL purified *Cj*PglB-biotin to a final volume of 30 μL and reactions were conducted in 1.5 mL microtubes in a 30°C incubator.

For on-chip glycosylation experiments, purified *Cj*PglB-biotin was immobilized on the functionalized surface of the device and the IVG reaction mixture was continuously pushed through the channels using a syringe pump with a total residence time of 30 min per chip. The reaction was heated by placing the microfluidic chip on a hot plate to maintain the internal temperature of the device at 30°C and confirmed by using a thermocouple in a similar arrangement. The product was then collected at the outlet of the device and either saved for analysis or recirculated through the device again to measure the effect of increasing residence times.

### 2.8 On-chip purification

The microfluidic device used for protein purification was designed with a series of posts at the end of the channel to entrap chromatography resin in the channel. For each device, we manually introduced Ni-charged profinity resin (Bio-Rad) into the channels before use. CFPS reactions expressing sfGFP^DQNAT-6xHis^ were then introduced into the inlet of the device with a total residence time of 30 min per chip and the outlet was collected and analyzed as the flowthrough fraction. The device was rinsed with PBS containing 10 mM imidazole at a flowrate of 2 µL/min and any unbound protein was collected and analyzed as the wash fraction. Finally, the target protein was eluted from the resin with PBS containing 300 mM imidazole at a flow rate of 2 µL/min. The fluorescence of each fraction was analyzed using a microplate reader to determine the GFP concentration and assayed for purity using standard SDS-PAGE with Coomassie blue staining.

### 2.9 Immunoblot analysis

For immunoblot analysis of IVG products and *Cj*PglB-biotin, samples were diluted 3:1 in 4× NuPAGE LDS sample buffer (Invitrogen) supplemented with 10% beta-mercaptoethanol (v/v). IVG products were boiled at 100°C for 10 min while *Cj*PglB-biotin samples were held at 65°C for 5 min. The treated samples were subjected to SDS-polyacrylamide gel electrophoresis on Bolt™ 12% Bis-Tris Plus Protein Gels (Invitrogen). The separated protein samples were then transferred to polyvinylidene difluoride (PVDF) membranes. Following transfer, the membranes containing IVG samples were blocked with 5% milk (w/v) in TBST (TBS, 0.1% Tween 20) and then probed with horseradish peroxidase (HRP) conjugated anti-His antibody (1: 5,000) (Abcam, catalog # ab1187) or rabbit polyclonal serum, hR6, that is specific for the *C. jejuni* heptasaccharide glycan (1:10,000) (kindly provided by Markus Aebi) for 1 h. To detect hR6 serum antibodies, goat anti-rabbit IgG conjugated to HRP (1:5,000) (Abcam, catalog # ab205718) was used as the secondary antibody. The membranes containing *Cj*PglB-biotin samples were blocked with 5% bovine serum albumin (BSA) (w/v) in TBST and then probed with ExtrAvidin-Peroxidase (1:10,000) (MilliporeSigma, catalog # E2886) for 1 h. After washing five times with TBST for 5 min, the membranes were visualized with Clarity ECL substrate (Bio-Rad) using a ChemiDoc™ MP Imaging System (Bio-Rad).

## 3 Results

### 3.1 Design of a modular microfluidic platform for continuous glycoprotein production

The design of our microfluidic-based glycoprotein biosynthesis platform integrated three key processes: protein expression, protein glycosylation, and protein purification (**Fig. 1**). In the first module of the device, sfGFP bearing a C-terminal DQNAT glycosylation motif^34^ that is optimally recognized by *Cj*PglB^37,42^ and a hexahistidine tag was expressed using crude S30 extract derived from *E. coli*, which enabled transcription and translation of the target protein on chip. We chose sfGFP as the acceptor protein so that the protein production and purification processes could be visualized and easily quantified during optimization of the microfluidic system. Next, in the second module, site-specific glycosylation was achieved by subjecting the newly expressed sfGFP^DQNAT-6xHis^ to components derived from a well-characterized bacterial *N*-linked glycosylation pathway, which occurs natively in the bacterium *C. jejuni* and has been functionally transferred to *E. coli*^36^. These components included *Cj*PglB as the glycan conjugating enzyme and *Cj*LLOs comprised of the *C. jejuni* GalNAc_5_(Glc)Bac heptasaccharide linked to undecaprenol-pyrophosphate as the glycan donor. *Cj*PglB and its cognate *N-*glycan structure were chosen here for proof-of-concept experiments because of the high transfer efficiency that has been observed with these components both *in vivo*^40,43^ and *in vitro*^18,24^. However, in a notable departure from previous works, we sought to site-specifically biotinylate *Cj*PglB and subsequently immobilize it in the device using biotin and streptavidin interactions, thereby enabling reuse of this important membrane protein biocatalyst^44^. Lastly, in the third module, the sfGFP^DQNAT-6xHis^ product was selectively enriched using a microfluidic device loaded with affinity resin that facilitated reversible capture of the hexahistidine-tagged glycoprotein product. The modularity of the device was designed to enable optimization of each unit operation and to allow flexible biosynthesis of different glycoproteins by simply interchanging acceptor protein target plasmids, glycosylation enzymes, LLO donors, affinity tags, and chromatography resins.

For the microfluidic device design, we aimed to create a system where protein synthesis, glycosylation, and protein purification could happen continuously in series at a fixed flow rate. Therefore, we fabricated individual chips to serve as building blocks that could be serially connected to increase the residence time of a particular process as needed. To test this design, we used an etched silicon wafer as a mold to fabricate channels in polydimethylsiloxane (PDMS) that was subsequently attached to glass slides. We chose PDMS because it enabled low-cost microfluidic fabrication that was sufficiently robust for device prototyping. Each microfluidic chip involved a serpentine channel design (width = 200 µm, depth = 120 µm, volume = 11 µL) that was inspired by previous work in which a similarly designed microfluidic bioreactor resulted in enhanced CFPS productivity^28^. Additionally, we hypothesized that long serpentine channels with a high surface area-to-volume ratio would promote efficient glycosylation by allowing sufficiently high levels of *Cj*PglB enzyme to be tethered to the device, thereby increasing the probability of contact with substrates. For the purification module, an immobilized metal affinity chromatography (IMAC) strategy was implemented whereby 25-µm posts were spaced apart from one another at the outlet of the device and the resulting channels were filled with Ni^2+^-charged beads for efficient hexahistidine-tagged protein capture.

### 3.2 Continuous-flow CFPS module improves protein production

As a first test of our design, we measured the on-chip protein titers obtained from the protein synthesis module following two modes of operation—batch and continuous flow—and compared these to the titers produced from one-pot reactions performed in standard microcentrifuge tubes. For these experiments, we generated crude S30 extract from *E. coli* strain BL21 Star™ (DE3) using a low-cost, sonication-based method ^41^ and the resulting extract was primed with plasmid pJL1-sfGFP^DQNAT-6xHis^ to drive the expression of sfGFP^DQNAT-6xHis^. In a standard 15-µL, one-pot CFPS reaction using a microcentrifuge tube, we produced 11.9 µg/mL of sfGFP^DQNAT-6xHis^ in two hours (**Fig. 2a** and **Supplementary Fig. 1**). To determine how the microfluidic environment affected sfGFP expression, we next performed batch-mode CFPS reactions in the first module of the microfluidic device. Specifically, the device was quickly filled with the same CFPS reaction mixture and fluorescence evolution was monitored in 30-min increments. When all CFPS components were present, fluorescence emission in the microfluidic channels gradually increased over time (**Fig. 2b**), corresponding to production of 9.4 µg/mL of sfGFP^DQNAT-6xHis^ in two hours (**Fig. 2a**). This result confirmed that the microfluidic environment itself had little-to-no effect on batch-mode CFPS productivity. It is also worth noting that surface blocking within the device was sufficient to allow sfGFP^DQNAT-6xHis^ clearance from the channels by simple rinsing.

**Figure 2.**
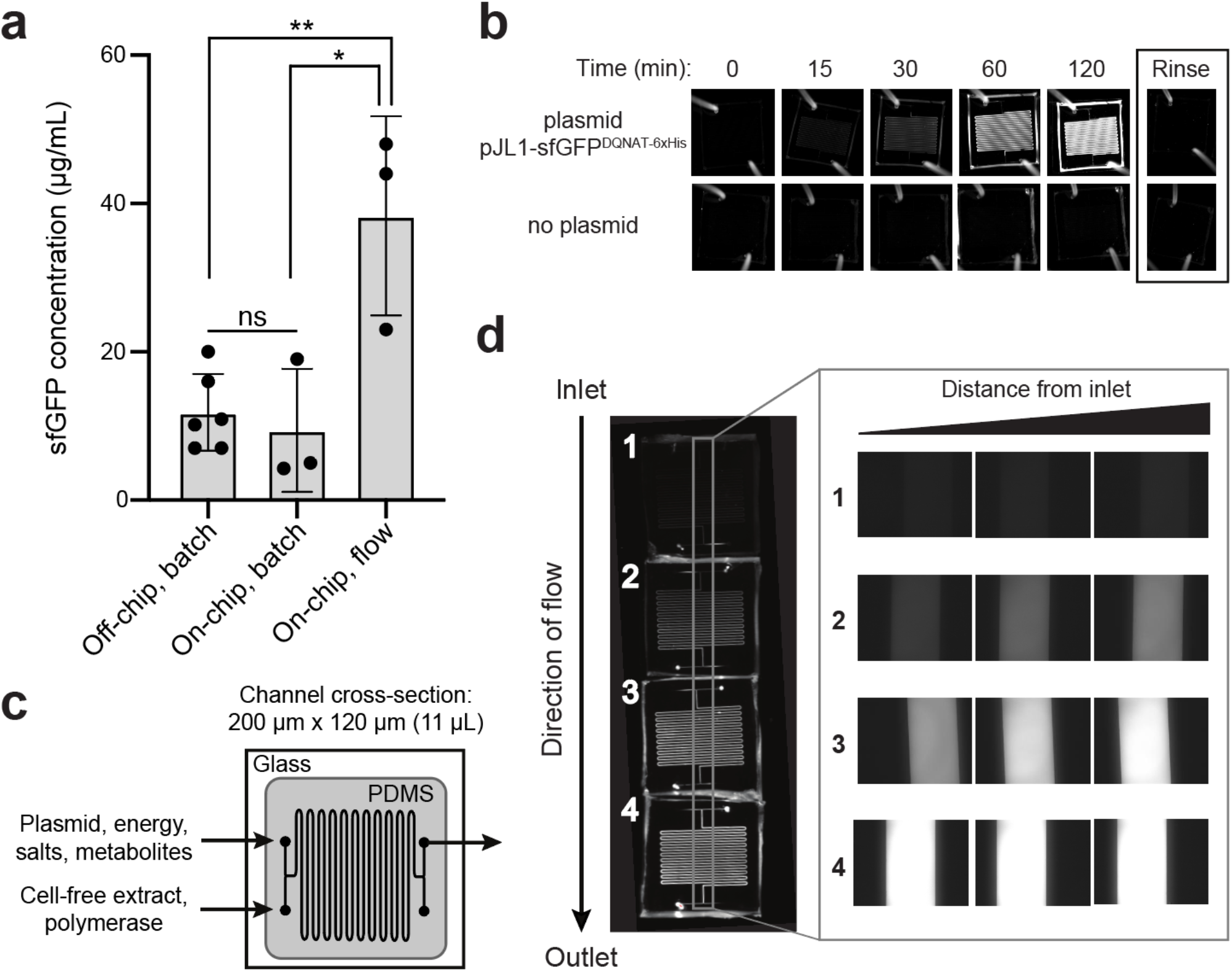
On-chip cell-free protein synthesis. **(a)** Fluorescence imaging of batch-mode operation in which all CFPS components were mixed, flown into the microfluidic chip, and allowed to react over a two-hour period. Representative images showing sfGFP^DQNAT-6xHis^ synthesis within the chip (top row) and a control experiment where plasmid was omitted from the CFPS reaction mixture (bottom row). **(b)** Mean sfGFP^DQNAT-6xHis^ concentration produced from the following reactions: off-chip, batch mode in a microcentrifuge tube; on-chip, batch mode in the microfluidic device; and on-chip, continuous-flow mode in the microfluidic device. For the on-chip systems, measurements were made on samples collected at the outlet of the chips. Data are the average of biological replicates (*n* = 3), error bars represent standard deviation, and *p* values were determined by paired sample *t*-test (*, *p* < 0.1; **, *p* < 0.01; and ns, not significant). **(c)** Serpentine channel microfluidic design for flow-based CFPS. The flow rate was set so that the reaction residence within each chip was 30 min. Cell-free extract and plasmid DNA were added at separate inlets so that protein synthesis was initiated inside the device. **(d)** Representative fluorescence images of continuous-flow mode in which four chips were linked together for a two-hour reaction residence time and reactants were flown into the two inlets. Inset shows expanded view of the regions within the gray box in the image at left.

We next investigated the effect of continuous flow on CFPS-based sfGFP expression. To generate a device that could accommodate a two-hour residence time (and thus be directly comparable to the batch-mode experiments above), we created a multi-chip system by linking individual devices with short pieces of tubing. Two input streams, one containing plasmid, energy, salts, and metabolites and the other containing S30 extract and T7 polymerase, met at the inlet and were mixed via diffusion between the two parallel streams as they moved through the channels (**Fig. 2c**). In a four-chip system, corresponding to a two-hour residence time, we observed increasing fluorescence along the length of the channels from the inlet to the outlet corresponding to production of 38.3 µg/mL of sfGFP^DQNAT-6xHis^(**Fig. 2a** and **d**). Fluorescence across the width of the channels was uniform, indicating that the solution was well-mixed. Additionally, when comparing the fluorescence generation in two-, three-, and four-chip systems, corresponding to one, one and a half-, and two-hour residence times, respectively, we observed non-linear protein production with the maximum production rate occurring between one and one and a half hours (**Supplementary Fig. 2a** and **b**). Importantly, the production rates in equivalent chips were similar, indicating that linking chips in series is a viable method for varying the residence time. Lastly, when comparing the titers of the two modes of on-chip operation relative to the microcentrifuge tube reaction for a two-hour residence time, we observed that sfGFP^DQNAT-6xHis^ produced on-chip in batch mode was statistically similar to off-chip production in a microcentrifuge tube, whereas introducing flow to the system significantly improved production by several-fold compared to both batch operations (**Fig. 2a**). This increase in production has also been observed by others^28^ and can be attributed to shorter diffusion lengths in the microfluidic channels.

### 3.3 Tethered OST enzyme enables a continuous-flow glycosylation module

In the protein glycosylation module, we sought to develop an OST tethering strategy that would allow for efficient protein glycosylation as the reaction substrates—the acceptor protein and LLOs— were continuously flown over the immobilized OST enzymes within the device. The advantage of OST tethering is that it enables creation of a local environment with a high concentration of OST enzyme that is reused in continuous operation. Such a reusable configuration is significant because OSTs are integral membrane proteins that are laborious and time consuming to produce in purified form^45^. For surface immobilization of *Cj*PglB, we leveraged avidin-biotin technology because it afforded the opportunity to site-specifically modify the OST with biotin such that enzymatic activity was minimally affected. To this end, an AviTag was genetically fused to the C-terminus of *Cj*PglB, providing a unique site for covalent biotin conjugation by separately prepared BirA enzyme. Biotinylation of *Cj*PglB was confirmed by immunoblot analysis using commercial ExtrAvidin-Peroxidase that specifically detects biotin (**Fig. 3a**). To verify that enzymatic activity of *Cj*PglB-biotin was not diminished by this modification or subsequent tethering onto a solid support, we performed off-chip *in vitro* glycosylation (IVG) reactions in a microcentrifuge tube using purified sfGFP^DQNAT-6xHis^ as acceptor protein, *Cj*LLOs as glycan donor, and either untethered *Cj*PglB-biotin or *Cj*PglB-biotin that was tethered to commercial streptavidin beads. Immunoblot analysis of the sfGFP^DQNAT-6xHis^ produced in these reactions was performed using an anti-His antibody to detect the protein/glycoprotein and hR6 serum that specifically recognizes the *C. jejuni* heptasaccharide. These blots revealed 100% conversion of sfGFP^DQNAT^ to the glycosylated form (g1) in reactions with both untethered and tethered *Cj*PglB-biotin, but only when the microcentrifuge tube for the latter reactions was shaken to keep the beads well suspended in solution (**Fig. 3b**). In the absence of shaking, the beads were observed to sink to the bottom of the microcentrifuge tube so that *Cj*PglB-biotin was not well dispersed within the reaction mixture, thereby reducing glycosylation efficiency as evidenced by the detection of sfGFP^DQNAT^ in a predominantly aglycosylated form (g0). Importantly, these results confirmed that *Cj*PglB tolerated both site-specific biotinylation and tethering to a solid surface without any measurable loss in enzyme activity.

**Figure 3.**
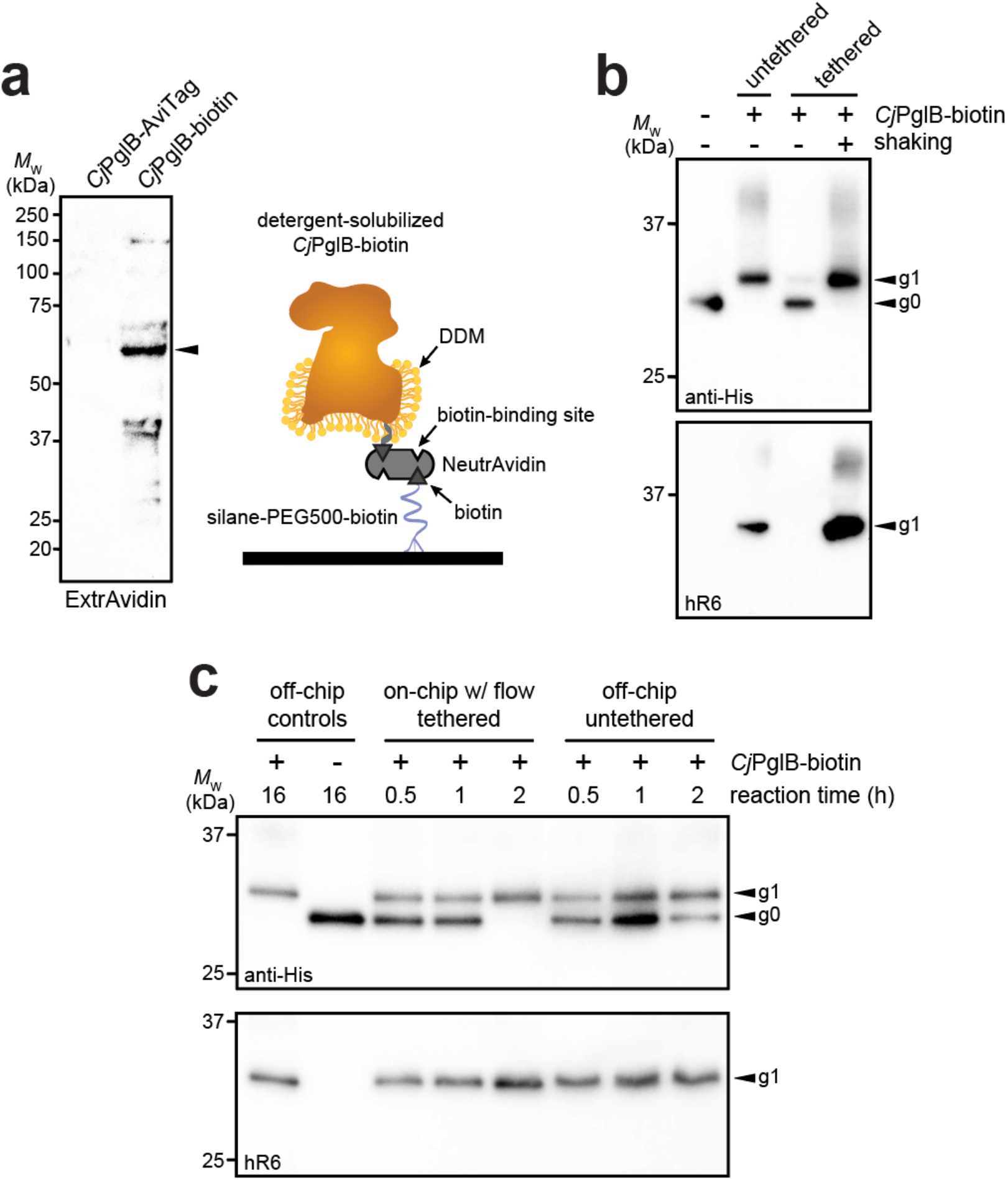
On-chip protein glycosylation. **(a)** Immunoblot analysis of *Cj*PglB bearing C-terminal AviTag that was subjected to biotinylation by treatment with BirA-containing lysate. Blot was probed with ExtrAvidin-Peroxidase that specifically detects biotin. Arrow denotes the expected molecular weight of *Cj*PglB-biotin. Schematic at right illustrates the tethering strategy used to immobilize *Cj*PglB-biotin generated in (a) within the channels of the microfluidic device. Schematic of *Cj*PglB-biotin tethering system. Silane-PEG5000-biotin was used to modify the surface of hydroxylated glass. Neutravidin, which has four binding sites with high affinity for biotin, was used to link *Cj*PglB-biotin to the surface of the device. **(b)** Immunoblot analysis of IVG reaction products generated in microcentrifuge tubes containing detergent-solubilized *Cj*PglB-biotin (untethered) or detergent-solubilized *Cj*PglB-biotin immobilized on streptavidin-coated beads (tethered). In the case of the latter, batch-mode reactions were performed with (+) or without (-) shaking as indicated. Blots were probed with an anti-polyhistidine (anti-His) antibody that recognized the C-terminal 6xHis tag on sfGFP^DQNAT-6xHis^ and hR6 serum that specifically recognizes the *C. jejuni* heptasaccharide glycan. **(c)** Immunoblot analysis of IVG reaction products generated using the on-chip, continuous-flow system with detergent-solubilized *Cj*PglB-biotin immobilized in the device channels (on-chip tethered) or the off-chip microcentrifuge system with detergent-solubilized *Cj*PglB-biotin free in solution (off-chip untethered). For the on-chip system, IVG components were flown through the channels, and the product was collected from the device outlet. Products from overnight microcentrifuge reactions in the presence (+) or absence (-) of *Cj*PglB-biotin were included as controls for glycosylation efficiency. Blots were probed identically as in (b). Arrows in (b) and (c) denote he monoglycosylated (g1) or aglycosylated (g0) sfGFP^DQNAT-6xHis^ products in each blot. Molecular weight (*M*_W_) markers are indicated at left of all blots.

Encouraged by these results, we went on to investigate a strategy for surface tethering of *Cj*PglB-biotin within the channels of our microfluidic device. To provide an evenly distributed, functionalized surface having low non-specific adsorption of other biomolecules, we modified the surface of our device with a silane-PEG5000-biotin moiety. This molecular weight of PEG has been shown to effectively reduce non-specific binding^46^ and to improve surface coverage compared to traditional coupling methods^47^. Here, silane-PEG5000-biotin provided a highly selective binding surface that was observed to promote higher loading capacity compared to non-specific adsorption to non-biotinylated silane-PEG5000 when visualized with fluorescently labeled streptavidin **(Supplementary Fig. 3a**). Comparing the surface coverage of the functionalized PEG brush to that of the non-covalent random adsorption also showed that we had a 30% increase in streptavidin coverage, allowing us to load more enzyme onto the surface of the device. Next, unlabeled NeutrAvidin was immobilized on the silane-PEG5000-biotin surface and was observed to bind fluorescently labeled, free biotin (**Supplementary Fig. 3b**), indicating that unliganded binding sites in surface-bound NeutrAvidin, which has four putative biotin-binding pockets, were available to capture additional biotin groups. Collectively, these experiments confirmed that silane-PEG5000-biotin provided a highly selective, passivating surface that increased binding capacity.

To evaluate this tethering strategy in the context of *Cj*PglB, we coated the channels of our microfluidic device with silane-PEG5000-biotin, followed by the addition of streptavidin and then *Cj*PglB-biotin (**Fig. 3a**). To determine whether immobilization of *Cj*PglB in this manner resulted in a glycosylation-competent device, we first performed on-chip IVG reactions in batch mode without flow. This involved manually pushing IVG reaction components—sfGFP^DQNAT-6xHis^ and *Cj*LLOs—over *Cj*PglB that was tethered in the microfluidic device. The sfGFP^DQNAT-6xHis^ product was collected from the chip and analyzed by immunoblotting, which revealed barely detectable glycosylation that was significantly less efficient than the glycosylation observed for an on-chip, batch-mode control reaction performed concurrently in a microcentrifuge tube (**Supplementary Fig. 4a**). To determine if continuous flow would remedy this issue, we next flowed the IVG reaction components over the device-tethered *Cj*PglB across a series of chips, each with a reaction residence time of 30 min. In parallel, batch reactions in microcentrifuge tubes were conducted at the same time for comparison. For these off-chip reactions, we calculated the maximum amount of enzyme that could theoretically be bound to the microfluidic surface and used that amount in the microcentrifuge-based reactions. It should be noted that this amount is likely higher than what is tethered within the device. The sfGFP^DQNAT-6xHis^ products from these reactions were analyzed by immunoblotting as above, with readily detectable glycosylation occurring in the on-chip, continuous-flow system that was on par in terms of efficiency with the off-chip microcentrifuge reactions (**Fig. 3b** and **Supplementary Fig. 4b**). Interestingly, the addition of flow even appeared to enhance the reaction kinetics, akin to what was observed in the CFPS module.

### 3.4 IMAC module enables continuous enrichment of product proteins

In the third module of our device, we sought to capture polyhistidine-tagged sfGFP^DQNAT-6xHis^ using an affinity capture strategy. By selectively binding our target protein, unwanted cellular debris, cofactors, and other waste products generated from the upstream reactions can be easily removed by flow-based rinsing. The glycoprotein product can then be recovered by elution with buffer containing a high concentration of imidazole. Using a design based on earlier works^31,48^, we prepared a PDMS microfluidic device with posts at the outlet that could be packed with commercial Ni^2+^-charged beads, thereby enabling on-chip IMAC (**Fig. 4a**). To test this strategy, we attempted to purify sfGFP^DQNAT-6xHis^in CFPS reaction mixtures that were flowed through the device with the initial exit stream collected as the flowthrough. Next, we switched the inlet stream to buffer for washing the IMAC resin and removing any non-specifically bound proteins. Finally, we eluted the hexahistidine-tagged protein product using imidazole. The loading and elution steps were monitored by fluorescence imaging of the device (**Supplementary Fig. 5a**) while the composition of each purification fraction was analyzed by SDS-PAGE analysis (**Fig. 4b** and **Supplementary Fig. 5b**). Based on multiple trials, we achieved 78 ± 10% purity in the final product (**Fig. 4b**). To determine the efficiency of product capture, we measured the fluorescence of each fraction and calculated the percent sfGFP^DQNAT-6xHis^ that was present. While there was some variation in the capture efficiency, we reproducibly captured 45 ± 14% of total produced sfGFP^DQNAT-6xHis^ (**Fig. 4c**). It should be noted that more complicated device configurations may improve the overall capture efficiency; nonetheless, our results are comparable to other microfluidic capture strategies^32^. This simple strategy for protein purification provides a convenient way to obtain a purified final protein product. Because of the modularity of our design, other types of resin (e.g., glycan-binding affinity reagents) could be used in place of, or in addition to the set-up shown here depending on the desired separation. Additionally, multiple devices could be connected for larger scale purifications.

**Figure 4.**
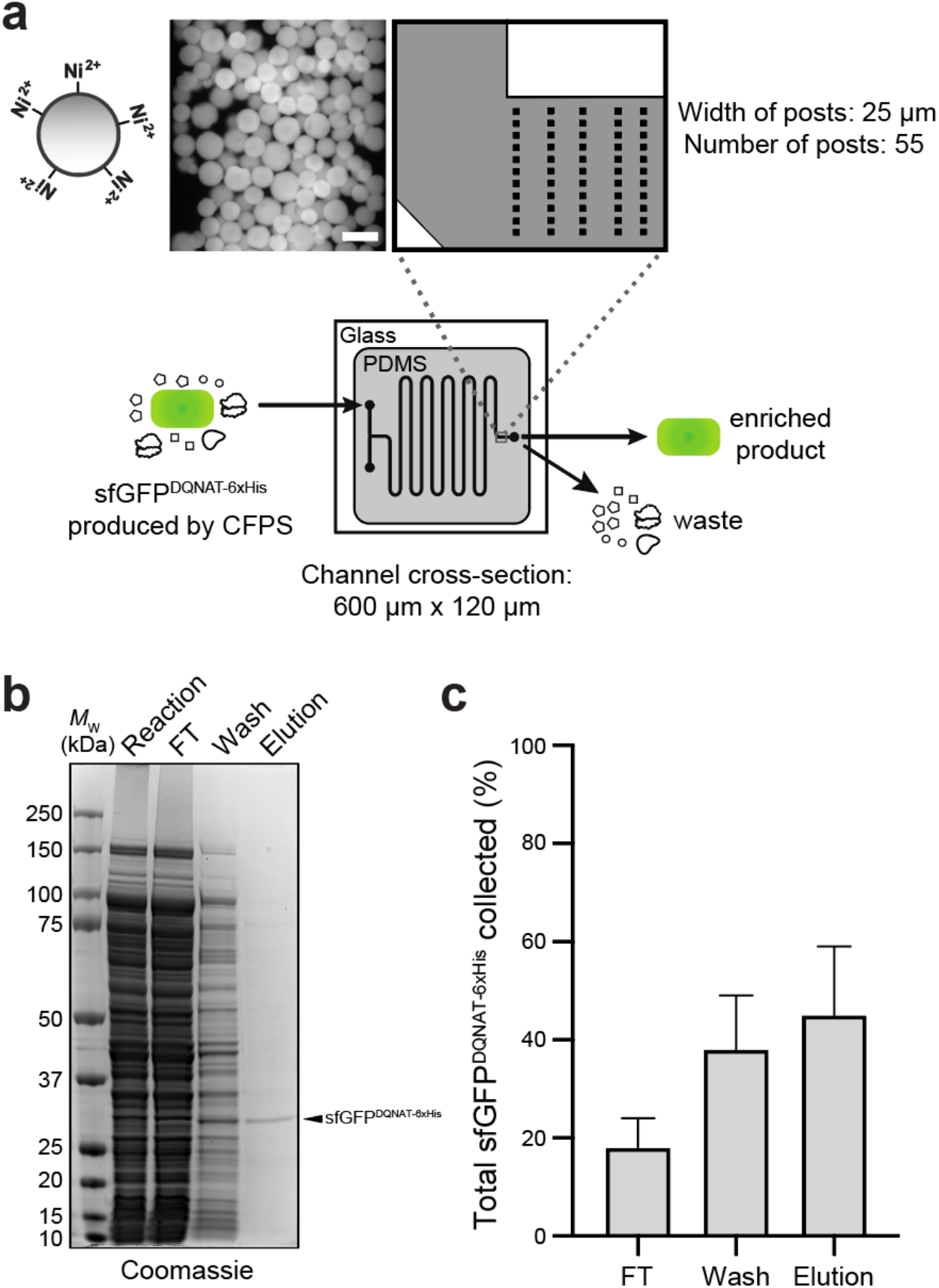
On-chip enrichment of CFPS product. **(a)** Schematic of purification module. The channels were designed to be 600-µm wide with fifty-five 25-µm posts at the outlet to accommodate Ni^2+^-functionalized beads. Shown at left is a representative fluorescence microscopy image of Ni^2+^-charged beads bound to sfGFP^DQNAT-6xHis^ within the device. After completion of CFPS reaction, product is pushed through the beads to allow for hexahistidine-tagged protein to bind to Ni^2+^ and flowthrough (FT) fraction is collected. Beads are then washed to remove any non-specifically bound proteins and wash fraction is collected. Finally, protein product is recovered through addition of buffer containing high concentration of imidazole and collected as elution fraction. **(b)** Representative Coomassie-stained SDS-PAGE gel comparing the protein composition of purification fractions as indicated. Arrow denotes the expected molecular weight of sfGFP^DQNAT-6xHis^. Molecular weight (*M*_W_) ladder is indicated at left. **(c)** Comparison of the amount of sfGFP^DQNAT-6xHis^ in each purification fraction represented as percentage of the total amount of sfGFP^DQNAT-6xHis^ collected. Data are the mean of biological replicates (*n* = 4) and error bars represent standard error of the mean.

## 4 Discussion

In this work, we designed and fabricated a microfluidic platform for flow-based, cell-free production of a model *N*-linked glycoprotein. This was accomplished in a modular system where protein synthesis, glycosylation, and purification were compartmentalized and individually optimized. In this approach, production rates were increased for continuous-flow processes compared to batch processes and protein production occurred at a faster rate than glycosylation. Importantly, the device was capable of glycosylating 100% of the added acceptor protein within two hours.

One of the most significant developments in this work was the demonstration that the pivotal glycosylation catalyst, *Cj*PglB, could be successfully immobilized within the device while maintaining high glycosylation efficiency. As a multi-pass transmembrane protein with regions in the membrane portion that are required for activity^49^, *Cj*PglB is challenging to express and purify; hence, the opportunity to reuse this enzyme in a continuous fashion should help to relieve a major bottleneck related to mechanistic studies of this enzyme and its biotechnological exploitation. Furthermore, the ability to achieve 100% glycosylation efficiency within the device allowed the glycoprotein product to be purified in a single step using IMAC. We anticipate that for less efficiently glycosylated proteins, an additional purification step using immobilized lectins or antibodies that specifically bind to the glycan could be implemented for glycoprotein enrichment.

For the proof-of-concept studies performed herein, we selected the *C. jejuni N-*linked glycosylation system as a model because of the flexibility of *Cj*PglB as a stand-alone, single-subunit OST^44^ that has proven to be compatible with a diverse array of glycan donors and acceptor protein substrates including some with therapeutic potential. To date, *Cj*PglB has been used to generate glycoproteins bearing bacterial^24,35,50^ and smaller human-type glycans^24,51–54^, and has enabled cell-free, one-pot systems for making *N-* and *O-*linked glycoproteins^24,55^ as well as antibacterial conjugate vaccines^56^. While not directly demonstrated in this work, cell-free strategies such as the glycosylation-on-a-chip platform described here could eventually provide access to glycoproteins that are modified with larger, complex-type *N-*glycans that mimic the structures commonly found on many human glycoprotein drugs such as monoclonal antibodies. This could be achieved by one-step *en bloc* transfer of fully assembled complex-type *N-*glycans or could instead be subdivided into discrete, compartmentalized modules. For example, we previously developed methods for *Cj*PglB-mediated transfer of the eukaryotic trimannosyl core *N*-glycan, mannose_3_-*N*-acetylglucosamine_2_ (Man_3_GlcNAc_2_), onto acceptor proteins both *in vivo* and *in vitro*^24,53^. The on-chip transfer of preassembled Man_3_GlcNAc_2_ glycans onto acceptor protein targets could serve as a first modular step that could be followed in subsequent modules by a series of immobilized GTs for elaborating the protein-linked Man_3_GlcNAc_2_ to discrete human-like *N-*glycan structures^57^. Alternatively, the ability of *Cj*PglB to transfer a single *N*-acetylglucosamine (GlcNAc) or diGlcNAc structure onto a target peptide^58^ provides a minimal glycan primer that could serve as an earlier starting point for single-enzyme transglycosylation using synthetic oligosaccharide oxazolines as donor substrates^59^ or multi-enzyme, cell-free glycan construction^23^. Importantly, our demonstration that *Cj*PglB can be immobilized in a microfluidic architecture without loss of catalytic activity is a critical first step to enabling any of these advanced strategies and paves the way for continuous production of a variety of therapeutically relevant glycoprotein products.

There has been increasing interest in the pharmaceutical industry to implement continuous manufacturing technologies that afford greater control over reaction variables, are amenable to automation, and are more flexible to changes in market demand compared to batch reactors^60–62^. Therefore, as a scaled-down model of flow-based systems, many researchers have investigated the use of microfluidic devices as microreactors for organic synthesis of pharmaceuticals^63,64^. Although biopharmaceuticals represent almost half of newly FDA approved therapeutics^65^, production of these more complex molecules by chemical means for incorporation into flow systems has been limited. Hence, our work expands the capability of microfluidic systems to now include production of *N*-linked glycoproteins by leveraging the open-box format of cell-free systems in a manner that provides spatiotemporal control over reactions, residence times, and concentrations. Looking forward, we anticipate that the flow-based glycoprotein production platform established here will inspire deeper exploration of cell-free technologies for continuous biomanufacturing of biologics.

## 5 Data Availability Statement

All data generated in this study are included in this article and its supplementary materials. Additional information is available from the authors upon reasonable request.

## 6 Author Contributions

A.K.A and Z.A.M are co-first authors of the manuscript and contributed equally to the experimental design, generation of data, and data analysis. Both have the right to list their name first in their CV, presentations, grants, etc. All authors contributed to project conceptualization, writing, and editing and have read and approved the final manuscript.

## 8 Acknowledgements

We thank Markus Aebi for providing strains CLM24 and SCM6 as well as hR6 serum used in this work. The authors also thank the Genomics Facility of the Biotechnology Resource Center at the Cornell Institute of Biotechnology for help with sequencing experiments and Thapakorn Jaroentomeechai, Han-Yuan Liu, Weston Kightlinger, and Mike Jewett for helpful discussions related to the manuscript.

