## Supplementary Information File for "Glycosylation-on-a-chip: a flow-based microfluidic system for cell-free glycoprotein biosynthesis"

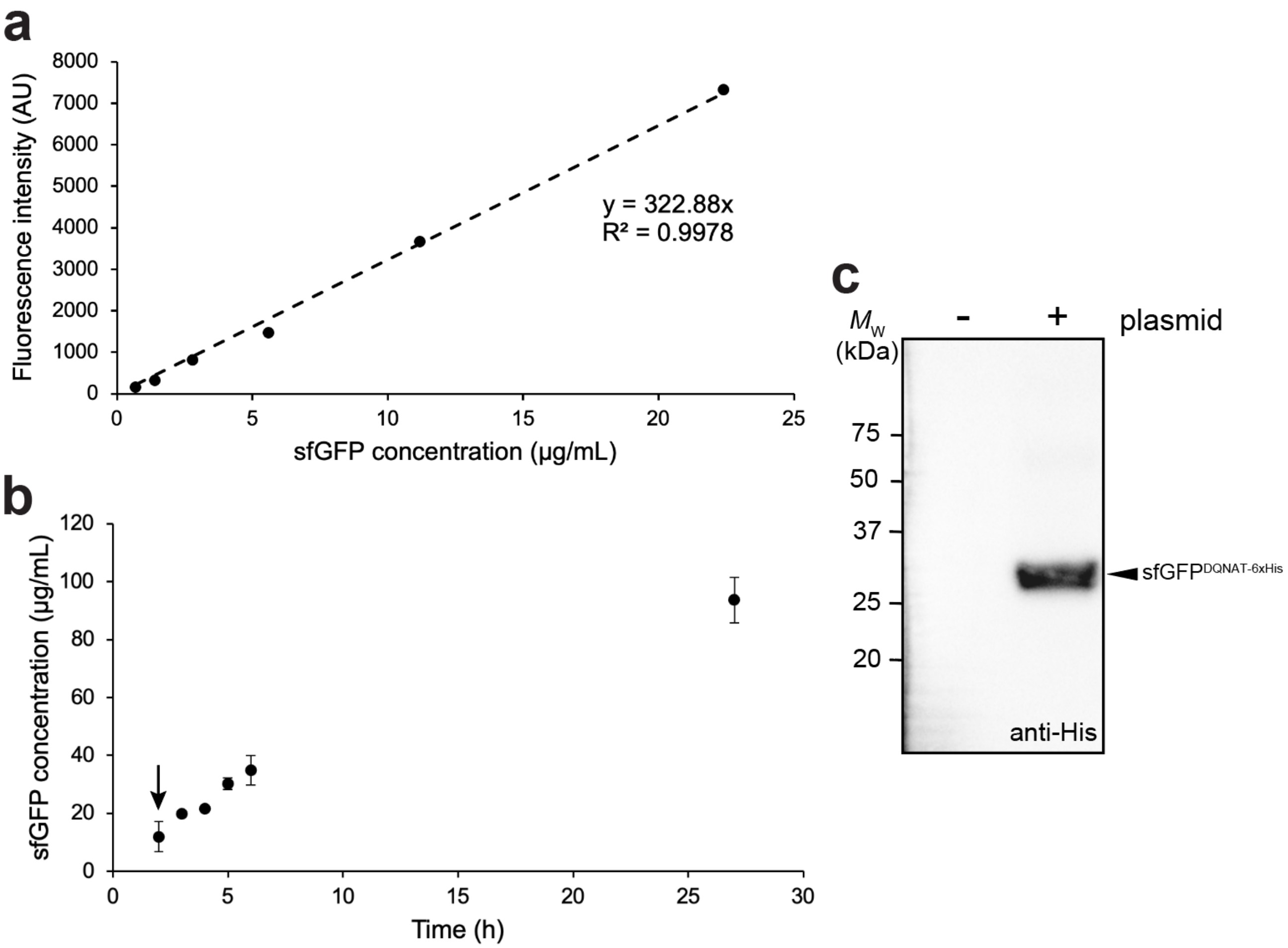


**Supplementary Figure 1. Characterization of sfGFP produced by CFPS. (a)** Calibration curve correlating sfGFP$^{DQNAT\text{-}6xHis}$ concentration to fluorescence intensity. Protein concentration was determined by Bradford assay while fluorescence intensity was measured on a plate reader at 485 nm excitation and 510 nm emission. **(b)** Effect of reaction time on batch-mode CFPS titers of sfGFP$^{DQNAT\text{-}6xHis}$. Cell-free extract was used to express sfGFP$^{DQNAT\text{-}6xHis}$ from plasmid pJL1-sfGFP$^{DQNAT\text{-}6xHis}$ and fluorescence intensity was monitored over time as in (a). The concentration of sfGFP$^{DQNAT\text{-}6xHis}$ produced in each reaction was determined by quantifying fluorescence intensity above background at different dilutions of the product to ensure that measurements were in the linear range of the calibration curve. Arrow denotes data point at $t$ = 2 h that is reported in Fig. 2a of the main manuscript. **(c)** Representative immunoblot of CFPS product using cell-free extract with (+) or without (-) addition of plasmid pJL1-sfGFP$^{DQNAT\text{-}6xHis}$ to the CFPS reaction mixture. Reactions were incubated overnight at 30°C and products were analyzed by immunoblot analysis in which membrane was probed with anti-polyhistidine (anti-His) antibody. Molecular weight ($M_W$) ladder is indicated at left of blot.

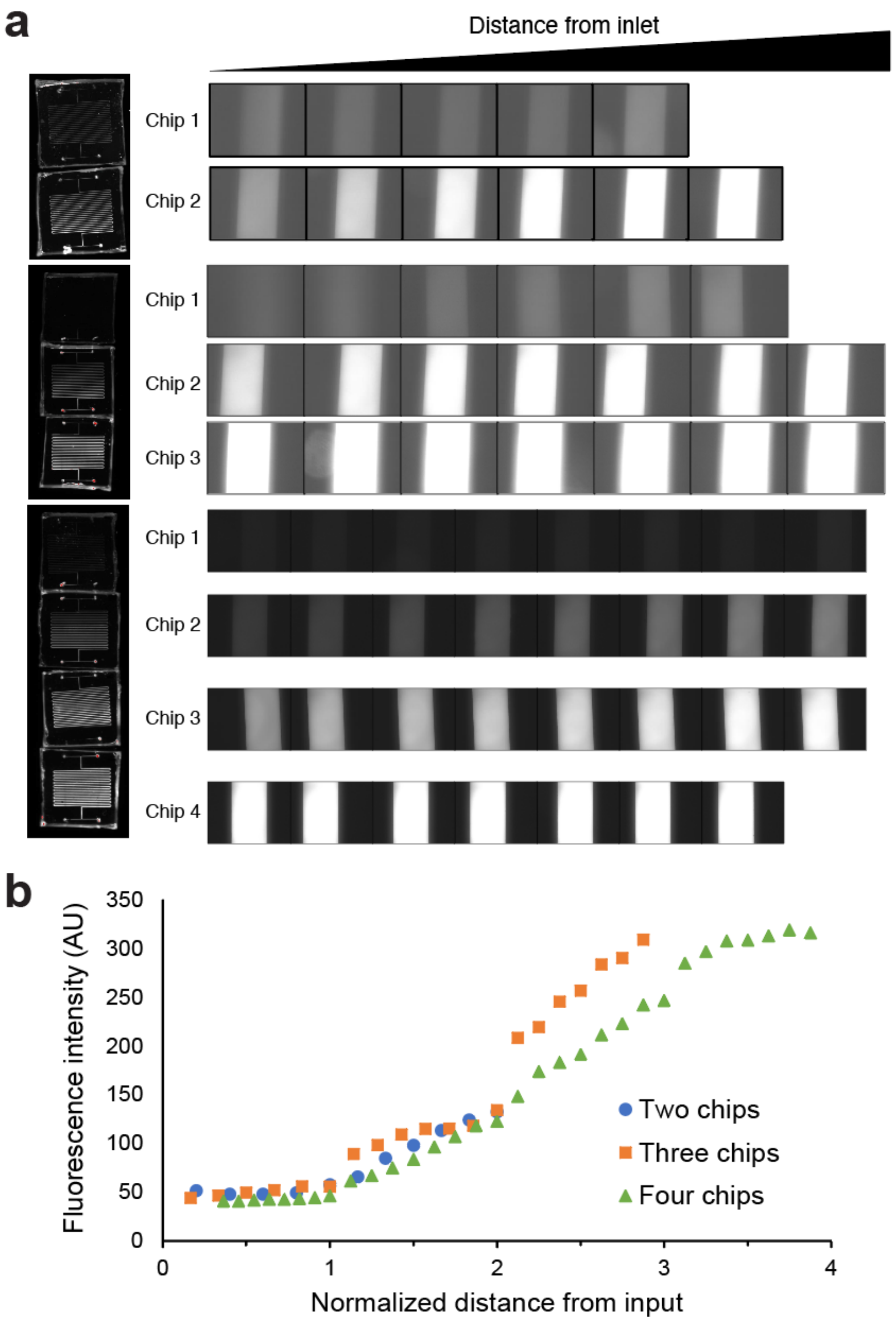


**Supplementary Figure 2. On-chip CFPS with continuous flow. (a)** In continuous-flow mode, two, three, and four chips were linked together for one-, one and a half-, and two-hour reaction residence times, respectively. Fluorescence evolution within the channels corresponding to sfGFP$^{DQNAT\text{-}6xHis}$ production by CFPS was monitored from the inlet to the outlet. Representative fluorescence imaging of the series of microfluidic devices were taken using a ChemiDoc imaging system with blue epi illumination. Images of the channels were taken with a microscope using a blue light source. **(b)** On-chip sfGFP$^{DQNAT\text{-}6xHis}$ production as determined by quantifying mean fluorescence intensity measurements from each microscope image in (a) plotted as a function of distance from the inlet of chip 1.

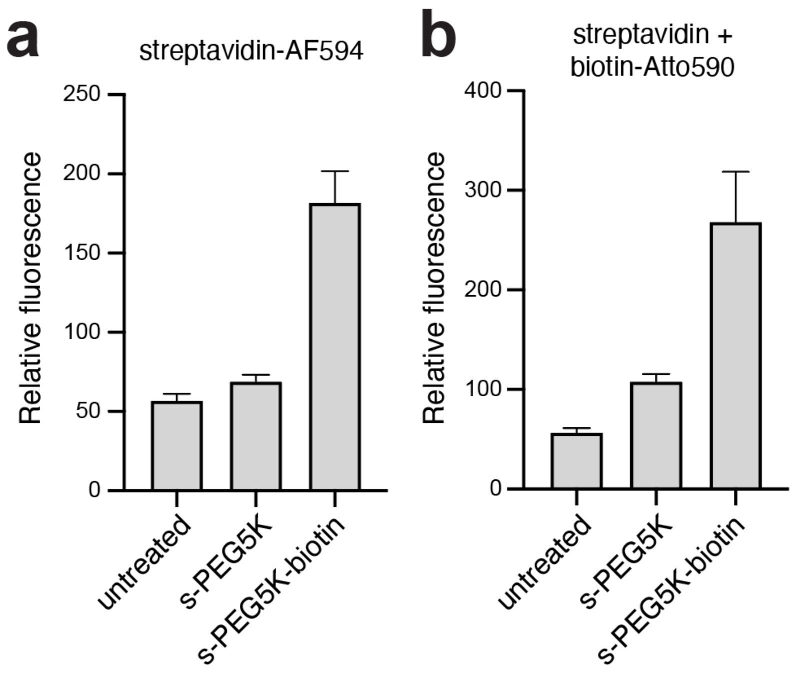


**Supplementary Figure 3. Protein immobilization on functionalized device surfaces. (a)** Fluorescence emission from microfluidic devices prepared with either silane-PEG5000 (s-PEG5K) or silane-PEG5000-biotin (s-PEG5K-biotin) following incubation with streptavidin conjugated with Alexa-Fluor594. An untreated device surface (untreated) subjected to the same amount of streptavidin conjugated with Alexa-Fluor594 served as a negative control. **(b)** Same treated and untreated microfluidic devices as in (a) but following a two-step treatment with non-fluorescently labeled streptavidin followed by biotin conjugated with Atto590. Fluorescence measurements in (a) and (b) represent the average and standard deviation of six biological replicates taken as a line scan across the width of the microfluidic channel. All data were normalized to fluorescence emission measured for untreated device.

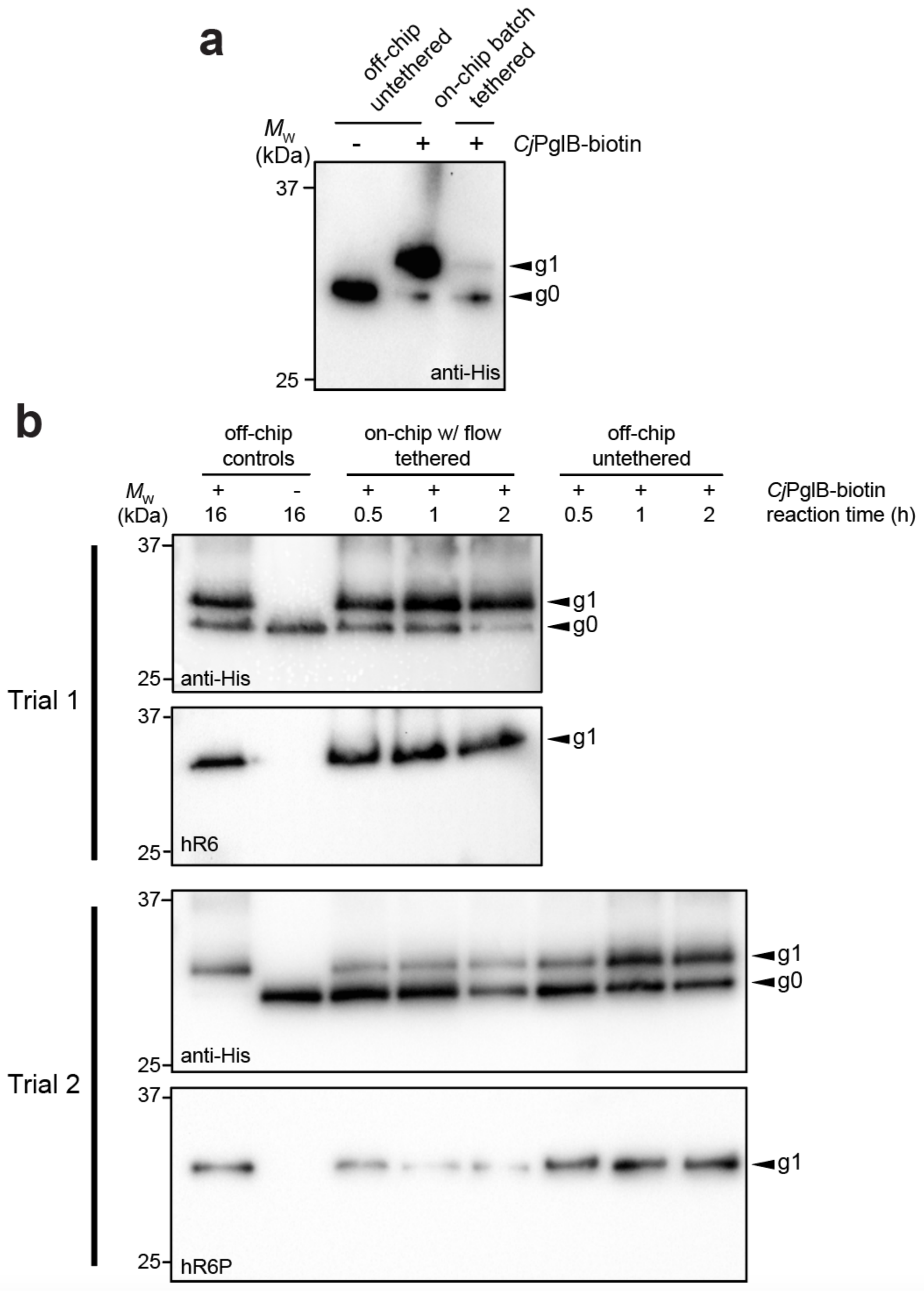


**Supplementary Figure 4. On- and off-chip glycosylation in batch and continuous-flow modes of operation. (a)** Immunoblot analysis of sfGFP$^{DQNAT\text{-}6xHis}$ glycosylation following either batch mode-mode IVG performed in microfluidic device with surface-tethered *Cj*PglB-biotin (on-chip batch, tethered) or in microcentrifuge tube with *Cj*PglB-biotin in solution (off-chip, untethered). For the latter mode of operation, a control reaction was performed without *Cj*PglB-biotin (-) as indicated. Reactions in all cases were carried out for 16 hours. **(b)** Biological replicates for on-chip continuous-flow mode experiments performed identically to those described in Figure 3c of the main manuscript.

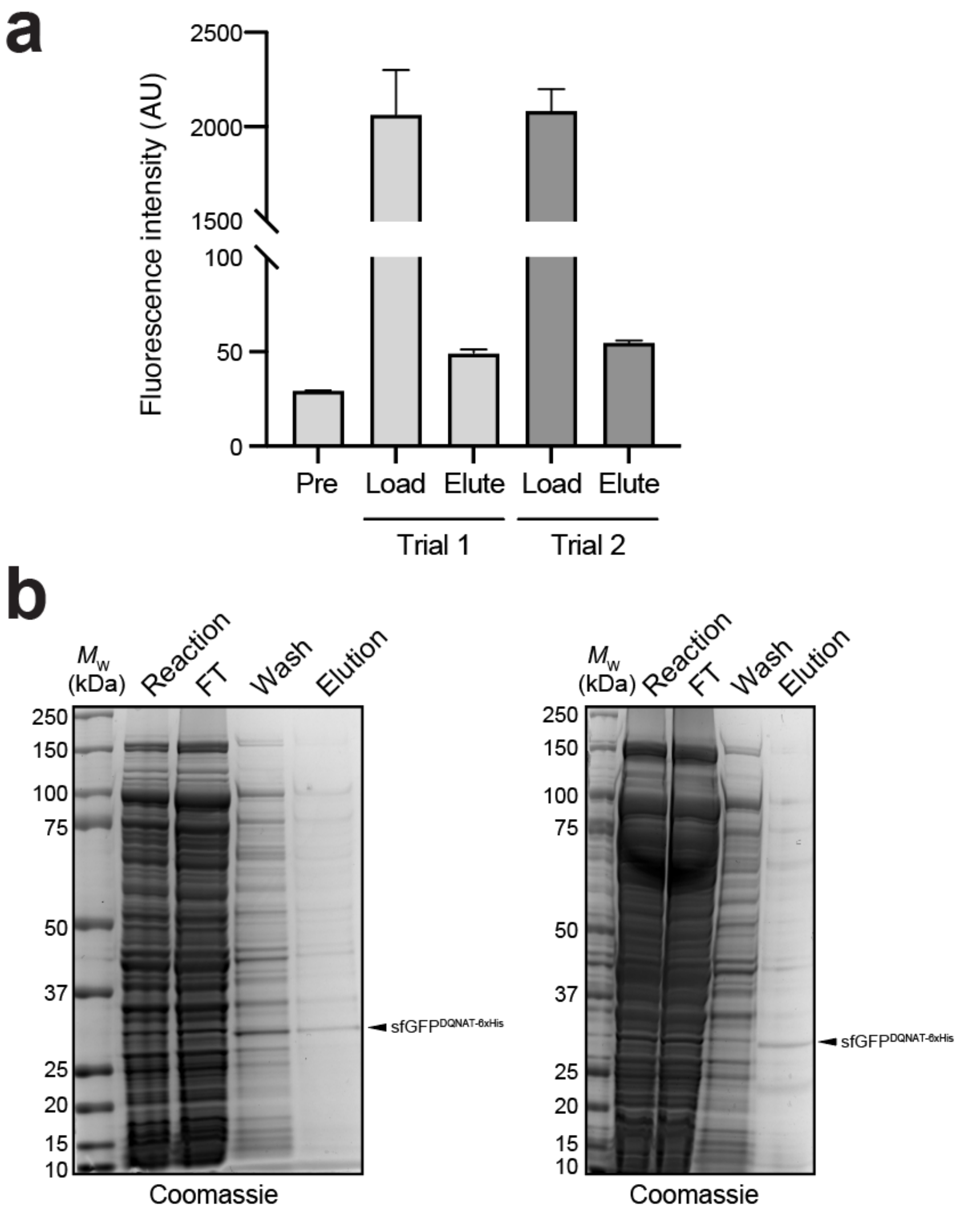


**Supplementary Figure 5. On-chip purification of glycosylated sfGFP$^{DQNAT-6xHis}$. (a)** Fluorescence imaging of purification module in which $Ni^{2+}$-charged beads were packed into the device channels and used to capture glycosylated sfGFP$^{DQNAT-6xHis}$. Background fluorescence within the channels was measured before sample loading (Pre) and compared to fluorescence after sfGFP$^{DQNAT-6xHis}$ was flown in the device and beads were thoroughly washed (Load) as well as fluorescence within the channels after 300 mM imidazole buffer was flown through the device (Elute). Trials 1 and 2 were performed subsequently using the same device, indicating that the purification module is reusable. **(b)** Biological replicates for on-chip purification experiments performed identically to those described in Figure 4b of the main manuscript. Arrow indicates the expected molecular weight of sfGFP$^{DQNAT-6xHis}$.
